# Impact of late larval nutritional stress on adult metabolic, gut and locomotor phenotypes in *Drosophila melanogaster*

**DOI:** 10.1101/2022.06.30.498321

**Authors:** Shri Gouri Patil, Sushmitha Sekhar, Aman Agarwal, TS Oviya, Debashis Rout, Megha

**Affiliations:** National Centre For Biological Sciences, Tata Institute for Fundamental Research, Bangalore, 560065 India; Centre For Ayurveda Biology and Holistic Nutrition, The University of Transdisciplinary Health Sciences and Technology, 74/2 Jarakabande Kaval, Attur Post via Yelahanka, Bangalore 560064 India

## Abstract

Dietary quantity and quality are key determinants for insect development from egg to adult. When nutritional deficiency is sub-optimal, development is completed, albeit resulting in an adult insect that is smaller than normal in size. If now fed a normal diet, would the smaller adults be similar to normally developed flies? To begin to answer this question, we characterised a few physiological and musculoskeletal readouts. Larvae were subject to acute starvation in late stages of development, and the resulting adults (Early Life Starved; ELS) maintained on a normal diet, were tested for biochemistry, gut physiology and locomotor activity. In females, no significant difference was observed in biochemical readouts for the whole-body or hemolymph, between control and ELS flies. In males, whole-body glucose and hemolymph trehalose were significantly reduced in ELS flies. Interestingly, ELS flies of both sexes respond with a disproportionally higher accumulation of triacylglycerides (TAGs) when on a high-fat diet. Age-related changes in the adult gut were compared between control and ELS flies: these revealed an increased proportion of ELS flies with loss of gut barrier integrity, deviant number of intestinal stem cells and no difference in enteroendocrine cells. The rate of antimicrobial peptide gene expression to an enteric infection challenge was also slower in ELS flies. For musculoskeletal readouts, climbing and flight behaviour were measured. In population assays, both male and female ELS flies showed climbing deficits. In a fine-scale climbing assay on individual flies, female but not male ELS flies showed higher climbing speed, while males but not females, showed lower geotactic index. This collection of phenotypic assessments show that firstly, larval undernutrition, even when not lethal, continues to impact adult functioning. Secondly, for some phenotypes, a normal diet in adults exposed to early life malnutrition is insufficient to restore optimal functioning. Thirdly, larval dietary loss affects adult insects in a sex-dependent manner. This study lays the framework to uncover the molecular, cellular and hormonal mechanisms that are altered by early life malnutrition. Furthermore, these readouts may be used to develop *Drosophila* as a high-throughput model for interrogating the efficacy of diet therapies to address malnutrition.

## Introduction

A fully formed and functional animal is the outcome of an exquisitely timed and organized developmental process; an activity that relies on several external and internal inputs. Food serves as an irreplaceable energy source that fuels this process and hence, both the quantity and quality of food matters greatly. Insect development is divided into four phases: egg, larva, pupa and adult. Of these, the larval and adult phases are inextricably dependent on externally available food sources. As *Drosophila melanogaster* gained popularity as a model animal, studies were undertaken to unravel the amount and type of nutrients that support fly development and maintenance (Begon *et al*., 1982). This has progressed into discoveries enabling us to outline molecular mechanisms that coordinate nutrition, development and metabolism (Droujinine and Perrimon, 2016). When viewed from the lens of life stages, such studies can be broadly classified into two types: 1) Understanding how nutrition and development are coordinated in larval stages or 2) interrogating how nutrition drives metabolism by studying adult responses. For instance, we know that fly development is sensitive to yeast concentrations (Slaidina *et al*., 2009; Okamoto and Nishimura, 2015) and that the larval fat body, coupled to systemic homeostasis via insulin and TOR signaling pathways, acts as an amino acid sensor to coordinate nutrient uptake and development (Layalle, Arquier and Leopold, 2008). In adults, feeding more yeast enhances fecundity and decreases lifespan (Partridge, Piper and Mair, 2005). High-sugar diets typically used to mimic diet-induced diabetes result in adults with high levels of circulating glucose and aberrant insulin signalling (Musselman *et al*., 2011). Not surprisingly, high sugar diets also result in a rapid onset of age-related decline in locomotory function and cardiac performance (Bazzell *et al*., 2013; Murashov *et al*., 2021). In larvae, high fat diets resulted in increased developmental time, accumulation of triacylglycerides (TAGs) and increased insulin signalling (Musselman and Kuhnlein, 2018). Adults on a high-fat diet not only show similar biochemical traits as larvae but also display decreased fecundity and lifespan, and increased cardiac defects (Birse *et al*., 2010). High-fat diets have also been shown to accelerate age-related behavioural senescence, such as reduced climbing ability (Liao, Amcoff and Nässel, 2021).

How larval nutrition affects adult behaviour and physiology, and its molecular underpinnings, is a relatively less explored area. When yeast concentrations are increased during larval stages, not only is developmental time reduced but also the final adult body size is proportionally bigger (Tu and Tatar, 2003). In the same study, larvae (juveniles) subjected to a low-yeast diet emerged as smaller size adults and when shifted to normal yeast diet in adulthood, displayed lower fecundity but normal lifespan (Tu and Tatar, 2003). In another study, higher yeast concentrations in the larval diet resulted in adults with higher expression of antimicrobial peptides, *Metchnikowin* and *Diptericin* in the female (Fellous and Lazzaro, 2010). On the other hand, high-sugar diets in larvae resulted in increased developmental time (Musselman *et al*., 2011) and as adults, such larvae displayed reprogrammed plasticity of sugar perception (May *et al*., 2019). In a recent chronic and switching experiment, wild-caught flies were fed variable yeast or sugar concentrations (2.5%, 10%, 25%) during development, and then switched into media of respective variable yeast/ sugar concentrations as adults. These adults were examined for fecundity, fat and glycogen content. Both low-yeast and high-sugar early life diets reduced fecundity, but only low-yeast diets resulted in increased fat content in adulthood (Klepsatel *et al*., 2020).

The studies reported above were designed to primarily change macronutrient ratios. Another way to study the nutrition and development paradigm is by keeping macronutrient ratios similar but reducing nutrient density by feeding larvae a diluted version of the respective laboratory’s standard fly food. This results in caloric insufficiency. Chronic larval nutritional deprivation (50% of normal diet) resulted in adult male flies with higher triglycerides and insulin signalling phenotypes (Rehman and Varghese, 2021). Another context for such investigations is the study of trade-offs in evolution. Chronic larval malnutrition (25% of normal diet) for several generations resulted in adults with poorer gut integrity upon infection (Vijendravarma *et al*., 2015), altered dietary nutrient absorption (Cavigliasso *et al*., 2020) and reduced host microbial diversity (Erkosar *et al*., 2018). In another set of experiments, adults selected for improved immune function over several generations and then exposed to chronic larval malnutrition (50% of normal diet), exhibited pathogen-specific susceptibility to infection (Singh *et al*., 2022).

We are motivated to model public health nutrition problems in *Drosophila melanogaster* and hence, chose an experimental system to model Developmental Origins of Adult Health and Disease (DoHAD). This concept arises from longitudinal cohort studies which observe that early life malnutrition not only impacts anthropometry but also greatly increases risk for cardiovascular diseases, metabolic dysfunction and mental health (Victora *et al*., 2008; Bhutta, 2013). In this article we outline the first set of phenotypic observations of adult flies subject to acute malnutrition in late-larval stages. As expected, nutritional deprivation leads to smaller adult size. What was surprising however, is that despite being on a normal diet post-eclosion, early life malnourished flies showed metabolic and behavioural phenotypes different from controls. Furthermore, the trend for differences between control and early life malnourished adults were not always in same direction in the two sexes. We also examined a few gut phenotypes in early life malnourished adults, and observed subtle differences in gut permeability and ability to deal with enteric infection. For behaviour, a difference in climbing and flight activity were observed. Together, these observations suggest that some adult phenotypes are more sensitive to nutritional insult during development.

## Materials and Methods

### Fly husbandry

A laboratory strain of *Canton S* was used in all experiments. Unless mentioned otherwise, flies were reared on “Normal Food”, 100mL of which consists of: 8g Corn meal, 4g Glucose, 2g Sucrose, 5g Yeast extract, 0.8g Agar, 0.4mL propionic acid, 0.7mL Benzoic acid, and 0.6mL of orthro-phosphoric acid. To generate ELS adults, 4hr egg lays were allowed to mature for 90 hours. L3 Larvae were subsequently transferred to either “Normal Food” (control) or 100mM (Sucrose) (Fig. 1A). After eclosion, adult flies were aged on normal food, at 25-30 adults per vial. Sexes were co-housed right up till experimentation. A brief exposure to ice was used to anaesthetize the flies, that were then sex-sorted for respective experiments. Consequently, all experiments have been performed on mated males and females. For high fat feeding, 5 day old flies were shifted to 10% D-Glucose(w/v), 4.5% active dry yeast (w/v), 7.6% agar (w/v), 20% virgin coconut oil (w/v), 0.1% methyl parahydroxy benzoate (dissolved in ethanol) (v/v) and 0.4% propionic acid (v/v) for 3 days. The control diet contained no coconut oil and 0.8% agar.

**Figure 1.**
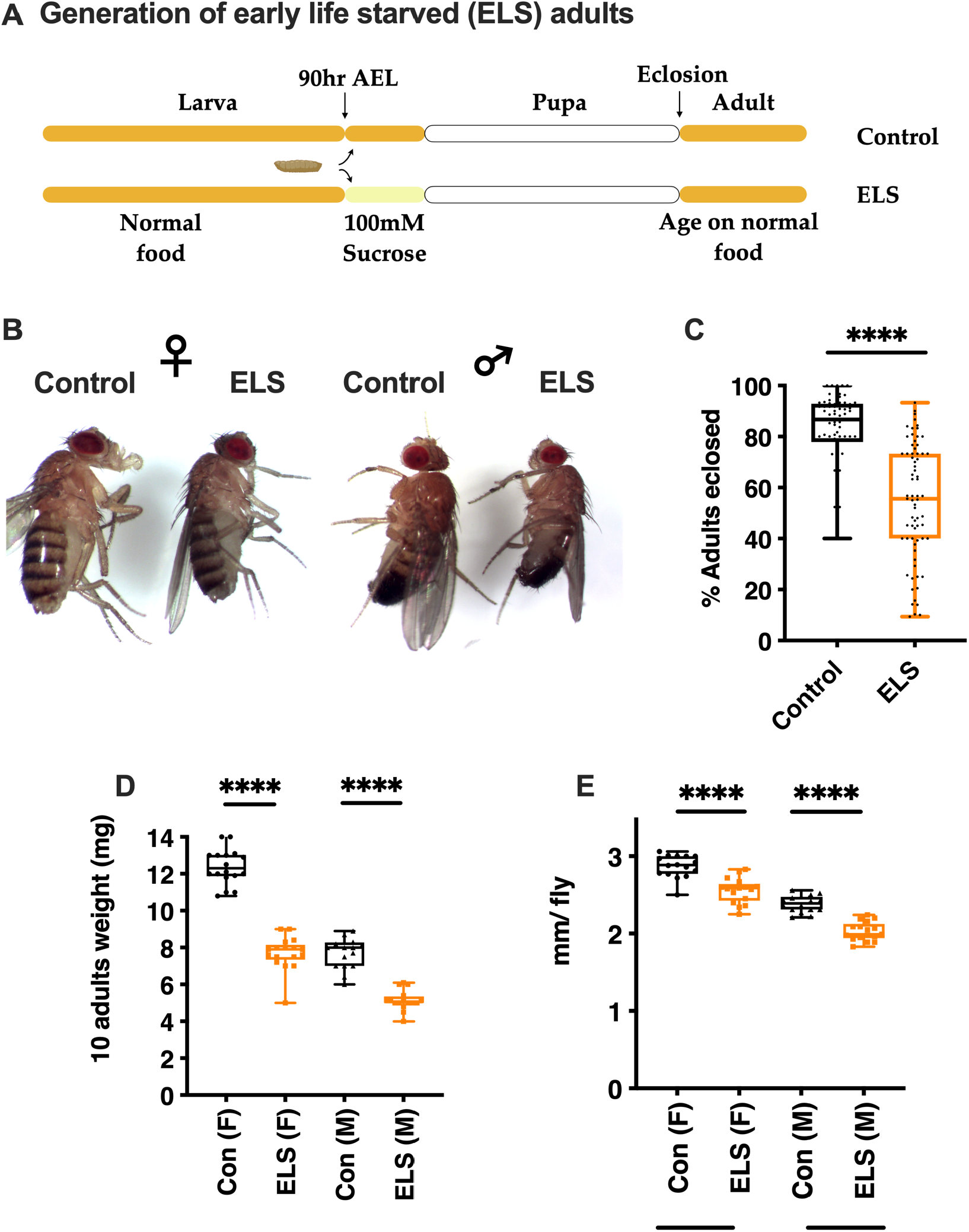
Anthropometry of adult flies subject to late larval nutritional stress. Protocol for making ELS adults **(A)** and representative images of eclosed adults **(B)**. Percentage of larvae that completed development to adulthood when placed on normal diet (control) or 100mM sucrose (ELS) n > 70 trials over a 1-year period. Each trial is a vial containing 30 larvae and monitored for total number of adults eclosed. unpaired *t*-test. ****p< 0.0001 **(C)**. Weight of 10 adults **(D)** sampled in different trials. n>15. unpaired *t*-test. ****p< 0.0001. Flies were anaesthetised and body length was measured as against a scale. n> 15, individuals sampled from different trials. unpaired *t*-test. p< 0.0001

### Metabolic assays

Flies were weighed in batches of 10 prior to homogenization. For protein and TAG, flies were homogenized in 0.05% Tween-20, heat inactivated at 70°C for 10mins, followed by centrifugation at 2200*g* at 4°C for 3mins. The resulting lysate was used to determine soluble protein using the BCA kit (Pierce) and TAG (Avantor). For glucose, trehalose and glycogen measurements, assay protocols as described by (Tennessen *et al*., 2014) were adapted. Trehalose is converted to glucose using trehalase, while glycogen is converted to glucose using aminoglycosidase. Standards as well as samples were incubated with respective enzymes for 22 hours. Finally, glucose with and without enzyme treatment was measured using the hexokinase reagent (Sigma; G3293). For lysates, flies were flash frozen and then homogenized in trehalase buffer (137 mM NaCl, 2.7 mM KCl, 5 mM Tris pH 6.6). After heat inactivation at 70°C for 10mins, followed by centrifugation at 2200*g* at 4°C for 3mins, the lysate was diluted in trehalase buffer and used either for trehalose and glucose detection or for glycogen and glucose detection. For glucose alone, lysates were prepared in trehalase buffer and immediately processed for detection. Concentrations were calculated against a standard curve generated by measurements of OD at 340nm. To collect hemolymph, flies were punctured in the thorax and 10 flies were placed in 0.5mL microfuge tube that had a hole at the bottom and was placed inside a 1.5mL microfuge tube. This set up was spun at 9000g, 4°C, 5 mins. 10μL of hemolymph collected at the bottom was diluted 1:100 with TB buffer and this diluted sample was used for protein, TAG, trehalose and glucose assays as described above.

### Gut related assays

Gut permeability was measured by feeding adults Eriogluacine (Sigma, 861146), a blue dye, as mentioned in (Rera, Clark and Walker, 2012). Flies were allowed to feed overnight and scored the next day. For immunohistochemistry, midguts were dissected in PBS (1.37mM NaCl, 27mM KCl, 100mM Na2HPO4, 20mM KH2PO4) and fixed with 4% PFA. Blocking was achieved with 5% Normal Goat Serum (NGS) and permeabilization by 0.5% Triton X-100 in PBS (PBS-Tx). Primary antibodies were freshly diluted to be used at 1:1000 (Mouse anti-prospero, DHSB, MR1A) and 1:2000 (Rabbit anti-pH3, Cell Signalling Technology, 9701) in PBS-Tx and applied for 24 hours. Secondary antibodies, Alexa Flour-488 anti-mouse IgG and Alexa Flour-568 anti-mouse IgG were at 1:1000, and applied for 1 hour. Before mounting, guts were stained with 1μg/mL DAPI for 30mins. Gut were mounted in 80% glycerol and imaged on Olympus BX41. ImageJ was used to count the number of enterocytes (DAPI stained cells), mitotically dividing stem cells (pH3^+^) and enteroendocrine cells (prospero^+^). Values are presented as proportion of positive cells to enterocytes, in a given area of the midgut. Only anterior (R4 and R5) regions were scored. RT-PCR was performed either on whole animals (3/sample) or dissected guts (5-6/ sample), but only in females. RNA was isolated using Trizol-Chloroform and cDNA synthesised as mentioned in (Megha and Hasan, 2017).

Primer pairs used are:

*rp49* CGGATCGATATGCTAAGCTGT, GCGCTTGTTCGATCCGTA

*Upd3* TGAACGAAACGCACAGCAAG; GTGATCCTGGCCTTGTCCTC

*Rel* ACAGGACCGCATATCG; GTGGGGTATTTCCGGC

*AttA* CCCGGAGTGAAGGATG; GTTGCTGTGCGTCAAG

*Dpt* GCTGCGCAATCGCTTCTACT; TGGTGGAGTGGGCTTCATG

For enteric infection, protocol described by (Buchon *et al*., 2010) was adopted. Briefly, an overnight stationary phase culture of *Erwinia cartovora cartovora* (Ecc15) was concentrated to OD_600_ of 200 and used to infect 10d old flies for the respective time. RT-PCR was performed on RNA extracted from whole female flies at the intended time points.

### Locomotor Assays

Climbing was measured by placing the flies in a glass cylinder. The apparatus was tapped gently and proportion of flies which were able to cross an 8cm mark in 12 secs were scored. Air puff stimulated tethered flight was measured as described in (Manjila and Hasan, 2018). Fine scale climbing assessment and analysis was performed as reported in (Aggarwal, Reichert and VijayRaghavan, 2019).

## Results

### Generation of adults (early life starved; ELS) subject to acute late larval dietary stress

Larval diet is an important determinant of final adult insect size. If larval diets are either of poor quality and/or of insufficient quantity there are two major consequences – an increase in developmental time and smaller adult size(Reviewed in Mirth and Shingleton, 2012). In *Drosophila melanogaster*, it is known that developmental time is refractory to food deprivation after larvae have achieved a body size called critical weight, ∼60 hours post-hatching (Mirth, Truman and Riddiford, 2009; Mirth and Shingleton, 2012). Hence, to understand the impact of acute nutritional loss, we designed a protocol where larvae, at the time well beyond critical weight (88-92hrs after egg laying) are moved to either 100mM Sucrose or normal food, and allowed to complete development, giving rise to undersized adults referred throughout as “Early Life Starved” (ELS) and control respectively (Fig. 1A, Representative flies: Fig. 1B). 100mM sucrose was used to mimic acute and severe malnutrition as it contains no protein or lipids or vitamins (typically obtained from yeast in normal food), approximately 50% less sugar and has been previously used to study nutritional stress during development (Megha and Hasan, 2017; Megha, Wegener and Hasan, 2019). From previous studies (Jayakumar *et al*., 2016; Megha and Hasan, 2017) we were aware that pupariation rates on this protocol for *CS* flies were high (> 90% on average). Here we report that it impacts pupal to adult transition significantly. On this protocol, on average, ∼ 83% of control and 54% of ELS adults were recovered (Fig. 1C). It suggests that nutritional insufficiency in the last 36 hours of larval development although refractory for pupariation, plays a significant role in pupal to adult development. The loss of approximately 36hours of acute nutritional stress is manifest in adult size, with flies that are ∼ 40% (females) and ∼30% (males) lower in weight, while being ∼11% (females) and ∼15% (males) smaller in length (Fig. 1D,E). Post eclosion, control and ELS adults were maintained at similar densities and sex-ratios on normal food (see materials and methods) till the indicated time for respective assays.

### Metabolic features of ELS adult flies

Nutritional inputs during development are experienced variably by different tissues, such that growth of imaginal discs and the brain are protected, while some like salivary glands and the adipose are not (Cheng 2012). The adult insect needs all organs to function normally and hence, to understand how cumulative health is affected, we measured readouts of gross physiology. No sex-specific differences between 5-6 day old ELS and control adults were observed for protein content (Fig. 2A). Because ELS flies are smaller in size, values reported for all systemic biochemistry parameters are normalized to weight. There are two energy storage molecules in the insect body – triacylglycerides (TAGs) and glycogen, which help the adult buffer nutritional stress. At steady state, ELS females showed a lower level of TAGs (Fig. 2B), and a modest increase in glycogen (Fig. 2C), but neither averages were statistically significant. As reported previously, levels of TAGs and glycogen are lower in males than females but between ELS and control males, no differences were observed (Fig. 2B and 2C). The predominant sugar molecule in insects is trehalose, although glucose also occurs in measurable amounts. No significant differences in trehalose and glucose were observed in control vs ELS females, but an interesting opposing trend was observed in ELS males. Although statistically not significant, ELS males tended towards higher trehalose levels and statistically, lower glucose levels.

**Figure 2.**
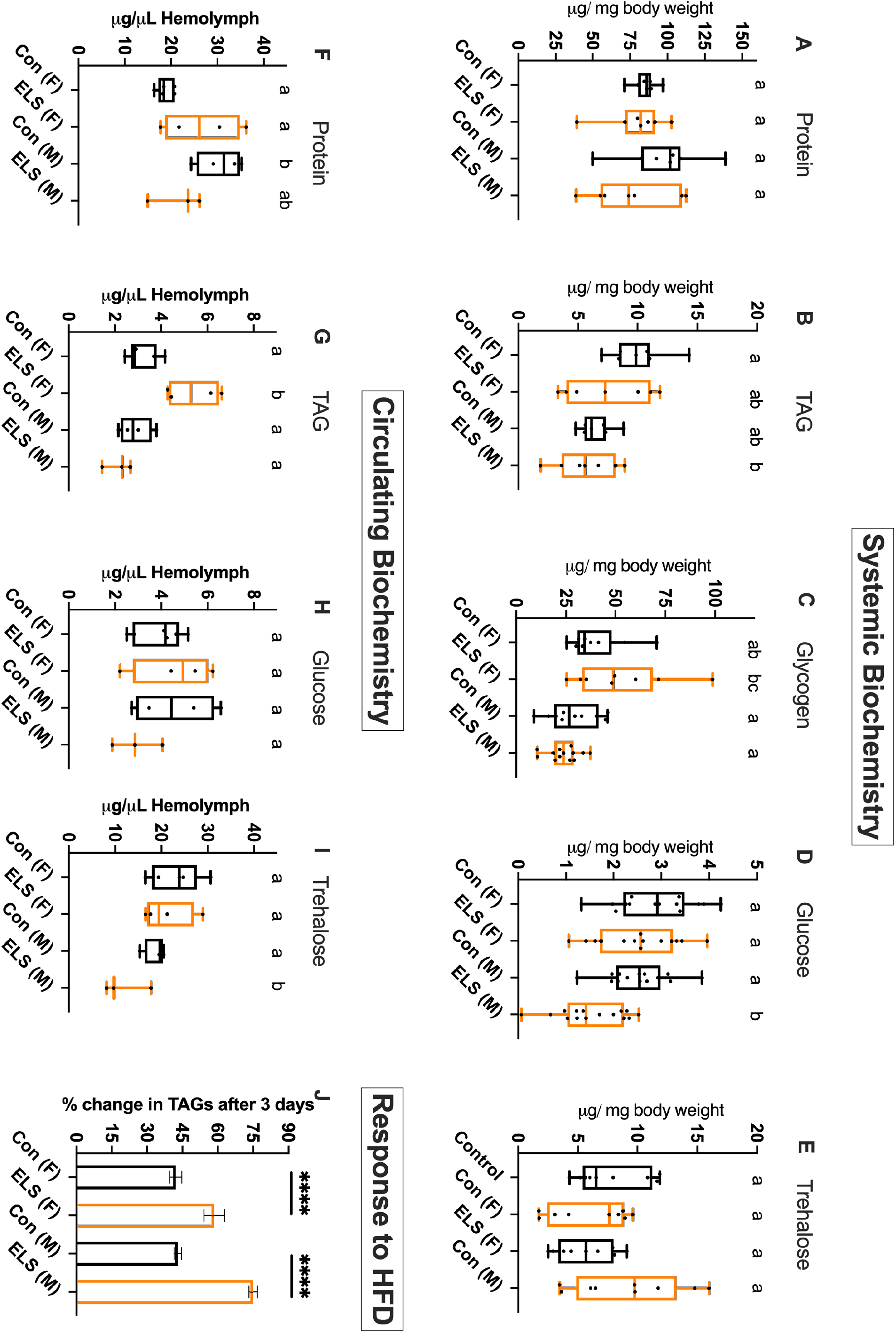
Biochemical parameters and response to high-fat diet (HFD). Whole adults were pooled, homogenised and biochemical assays performed as mentioned in Materials and Methods for protein **(A)** n=7, triacylglycerides or TAGs n=7 **(B)**, glycogen n>8 **(C)**, glucose n>14 **(D)** and trehalose n = 9 **(E)**. Hemolymph was collected from 10 flies and biochemical assays performed as mentioned in Materials and Methods for protein **(F)**, TAGs **(G)**, glucose **(H)** and trehalose **(I)**. n=4 except for ELS males where n=3. Ordinary one-way ANOVA with Tukey’s multiple comparison test. Bars with the same alphabet represent statistically indistinguishable groups. 7 day old flies were placed on a diet with or without 20% coconut oil (high-fat diet) and TAGs were measured after 3 days. % change in TAGs in flies on HFD as compared to control diet is reported. n=6. unpaired *t*-test performed separately for females and males. p< 0.0001

To further understand if there are homeostatic imbalances in biochemistry, nutrient levels were measured in the hemolymph. Interestingly, ELS females, like control males, displayed higher levels of protein as compared to control females (Fig. 2F). However, protein levels in ELS males are lower than control males (Fig. 2F). For TAGs, circulating levels are highest for ELS females and in light of their lower systemic TAG levels (Fig 2B), it suggests that lipid homeostatic mechanisms are differently regulated in ELS females. Circulating levels of glucose were not different in either sexes or conditions (Fig 2H), but levels of trehalose were lower in ELS males as compared to control males (Fig. 2I). Trehalose is particularly interesting as on an average, it was higher in whole body lysates of ELS males (Fig. 2E). The glucose and trehalose data (Fig. 2D,E,H,I) suggest that sugar metabolism in ELS males may be perturbed.

Together, the systemic and circulating biochemistry of ELS adults reveals some interesting variations based on sex and nutrients. It must be noted here that these differences persisted even after adults were on normal food for 5-6 days. Further studies on specific nutrient pathways are required to understand how these differences lead to functional changes. Here, we tested one such functional change i.e., response to high fat diet. Epidemiological studies in humans, have shown a strong correlation between low birth weight and high adult risk for cardiovascular disease and metabolic health. Hence, we wondered if a similar risk is observed in our model. Control and ELS adults were subject to 3 days of a high-fat diet and the level of TAGs measured. This protocol is known to increase TAG levels in flies (Birse *et al*., 2010). Remarkably, in both male and female ELS adults, the amount of TAG accumulated is more than in controls (Fig. 2J). This suggests that a sex-independent program that buffers metabolism against excess fat consumption is altered in ELS flies.

### Gut Parameters

A syndrome called Environmental Enteric Dysfunction (EED) is reported in malnourished children from low and middle income countries (Crane, Jones and Berkley, 2015). The cause of EED is unknown and the symptoms are a collection of various secondary readouts for inflammation as well as gut function. One hypothesis for EED is that the post-natal gut is still developing and repeated assault by enteric infections in the context of undernutrition results in increased gut permeability and higher levels of inflammatory molecules.

To understand if EED can be modelled in *Drosophila* we first sought to understand if gut health is altered in ELS adults. Gut permeability was measured using a popular dye feeding assay that classifies flies as “smurfs” if the blue dye is spread across the animal, thus reflecting high gut permeability as opposed to “non-smurfs” which retain the blue dye in their gut (Rera, Clark and Walker, 2012). In control flies, as expected, the proportion of smurfs increases with age. However, irrespective of age, a small but significantly higher proportion of ELS flies were found to be smurfs indicating a higher number of animals with compromised gut permeability (Fig. 3A). A leaky gut is also associated with infiltration of immune cells and higher inflammation, hence we measured the gene expression of *Drosophila* cytokine homologues of TNFα (*eiger*) and IL-6 (*upd3*), as well as a transcriptional factor that regulates antimicrobial peptide production, *relish (rel)*. At least on day 10, neither in the midgut nor in whole body, was the expression of these genes elevated (Fig. 3B).

**Figure 3.**
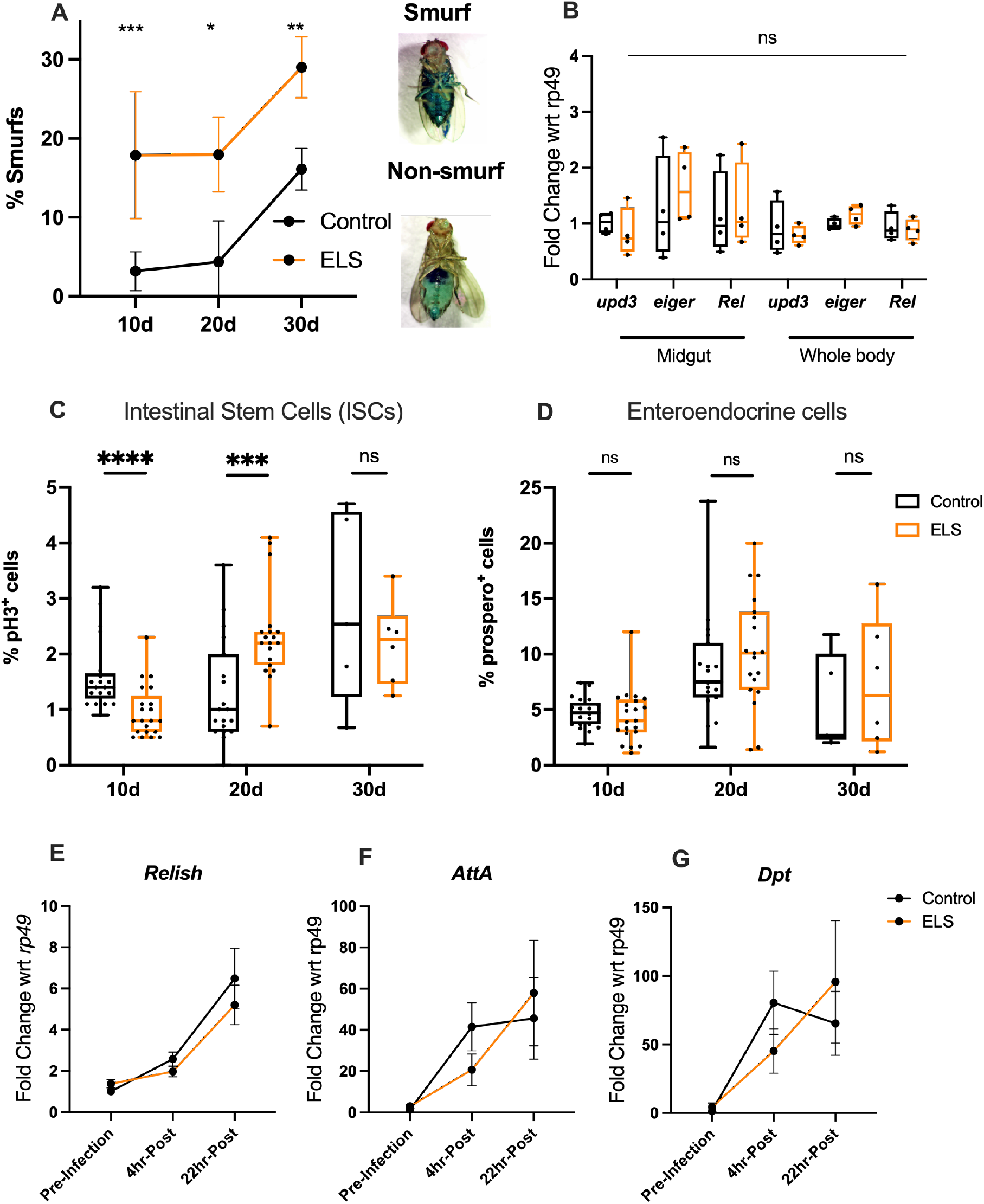
Gut-related investigations. **(A)** Proportion of flies with poor gut barrier integrity (smurfs) over time. n=5 trials of 30 flies each. Mann-Whitney test. *** p< 0.001, ** p< 0.01, *p<0.05. Ranks compared by Homs-Sidak method. RT-PCR was used to measure fold change in gene expression of inflammatory signals *upd3, eiger* and *Rel* (*relish*) either in dissected midgut or whole body **(B)**. Dissected midguts were stained with anti-phospho-histone (pH3) or anti-prospero antibodies to mark mitotically dividing cells (ISCs) **(C)** or enteroendocrine cells **(D**) respectively. Only cells in the R4 and R5 regions of the midgut were counted. % positive cells were normalised to total number of enterocytes in the observed area. Mann-Whitney test. **** p< 0.0001, *** p< 0.001. Ranks compared by Homs-Sidak method. n > 18 for 10 d (day), 20d -old flies while n = 6 for 30d. **(E-G)** RT-PCR was used to measure fold change in gene expression of transcription factor *relish* (rel) and downstream AMPs, *attacin* (*AttA*) and *diptericin* (*Dpt*) in whole body lysates of flies infected with *Ecc15*. n=8.

The *Drosophila* midgut consists of four major cell types: intestinal stem cells which give rise to all other cell types, an intermediate enteroblast cell on its way to becoming either an absorptive enterocyte or hormonal enteroendocrine cell (Miguel-Aliaga, Jasper and Lemaitre, 2018). Intestinal stem cells divide regularly and the dividing cells can be marked by the phosphohistone antibody (pH3^+^), while enteroendocrine cells are marked by staining with *prospero*, a transcriptional factor (Ohlstein and Spradling, 2006). At day 10 and day 20, the proportion of ISCs normalized to the total number of enterocytes was observed to be different in ELS vs control flies (Fig. 3C). This difference disappears at day 30 (Fig. 3C). No significant difference was observed in the number of enteroendocrine cells (Fig. 3D).

To further characterize gut health we monitored the innate immune response of day 10 adult flies to a non-lethal enteric infection by *Erwinia carotorova carotorova* (*Ecc15*). Infection by a gram negative bacteria activates the IMD pathway in *Drosophila*, leading to the increased expression of a transcriptional regulator Relish (*rel)*, which in turn drives the expression of several antimicrobial peptides (AMPs) genes of which two, AttacinA (*AttA*) and diptericin (*Dpt*), were measured here (Bruno, Jean-Marc and A., 1997). Interestingly, while the up-regulation of *Rel* was similar between ELS and control adults, the rate of up-regulation of *AttA* and *Dpt* was slower in ELS flies (Fig. 3G). 4hr post-infection, control flies exhibit close to their maximal up-regulation of AMPs, while ELS adults are only at half-maximal expression (Fig. 3G). By 22 hours however, ELS adults are expressing more AMPs than control flies (Fig. 3F, G). It must be noted here that these differences were not statistically significant.

### Locomotor assays

While doing routine transfers between vials during ageing, we observed that ELS flies when dropped into the vial, would take longer to right themselves and climb the walls of the vial. Therefore, we decided to investigate if their locomotory behaviours were affected. In a simple cylinder climbing assay, both ELS males as well as females displayed a climbing deficit (Fig. 4A). While adult flies show decreased locomotor activity with age, it was unexpected to find this deficit in 5 day old ELS flies. Next, we measured air-puff stimulated tethered flight in these adults and again, observed differences between ELS and control adults (Fig. 4A). Here however, males and females show opposing trends – female ELS adult show less flying time than controls, while ELS males show higher flying time (Fig. 4B). This behaviour is not immediately consistent with observations on systemic or circulating biochemistry (Fig 2A-I). For e.g., lower levels of glycogen, an energy storage molecule, might be expected to correlate with lower flying time (as seen in control females vs males). In ELS adults, females have comparatively higher glycogen levels (Fig. 2C) but lower flight times, and ELS males have lower glycogen levels but higher flight times.

**Figure 4.**
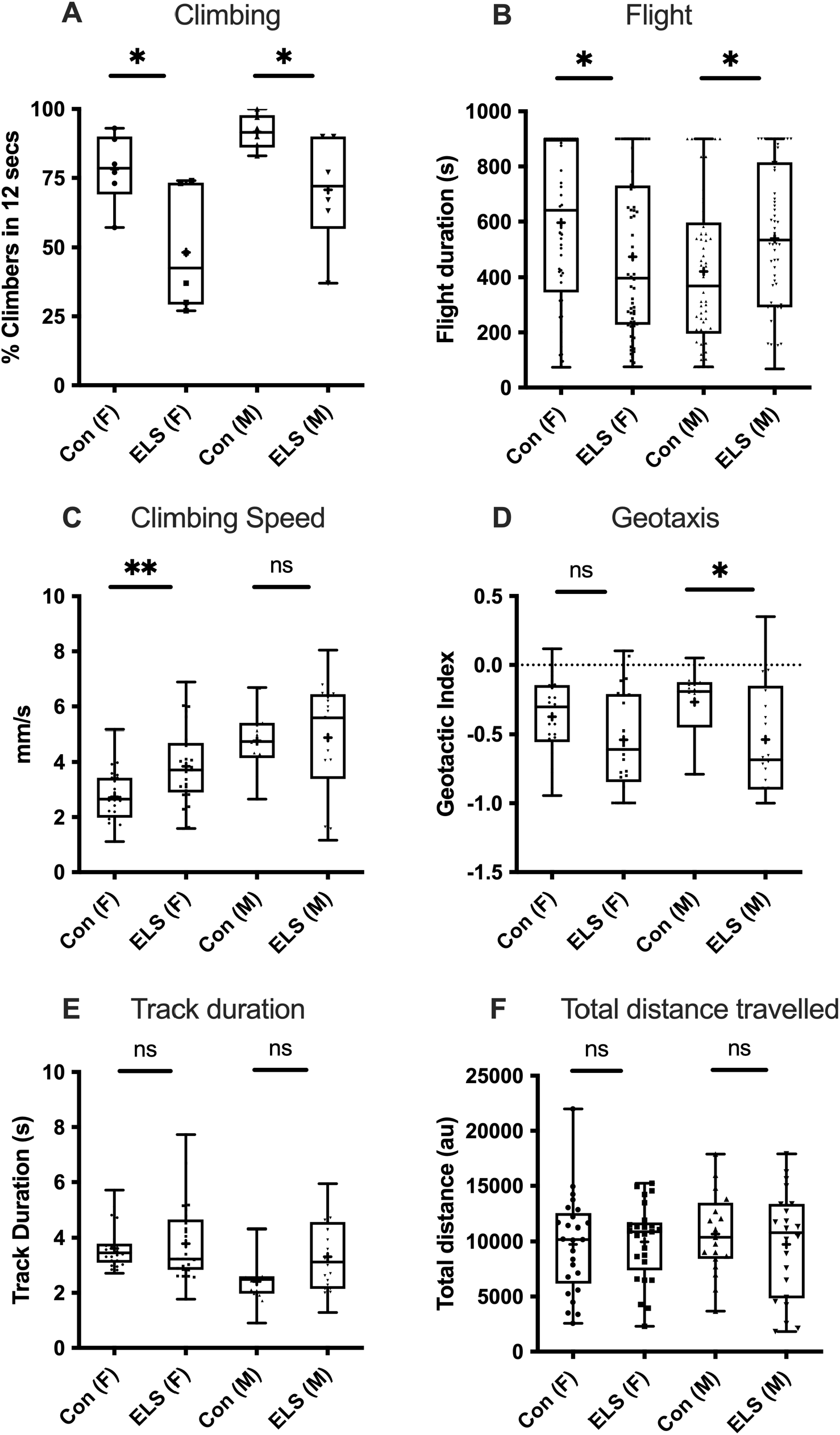
Locomotory phenotypes. **(A)** Climbing assay. n=6 batches of 10 flies each. Unpaired t-test. * p < 0.05. **(B)** Air-puff stimulated tethered flight. n > 59 flies. Mann-Whitney test. * p< 0.05. Ranks compared by Homs-Sidak method. Fine-scale climbing assessments of individual flies for speed **(C)**, geotaxis **(D)**, track duration **(E)** and total distance travelled **(D)**. two-tailed unpaired t-test. ** p< 0.01, * p < 0.05. n>19.

The climbing and flight assays provide population-level behavioural information. To augment these with individual data, we deployed a fully automated climbing assay that characterizes multiple locomotory features at fine-scale. As reported previously (Aggarwal, Reichert and VijayRaghavan, 2019), climbing speed of males is higher than that of females (Fig. 4C). Interestingly, ELS males and females showed a higher climbing speed, although only the average climbing speed of females was statistically significant (Fig. 4C). Combined with the climbing deficit data (Fig. 4A), it suggests that ELS flies are unable to cover a long distance, but the distance they do climb, they do so faster. When climbing, flies are negatively geotactic i.e., move away from gravity. Both ELS females and males were observed to have a lower geotactic index (Fig. 4D), suggesting issues with their ability to either sense gravity or this could be a reflection of their poor climbing ability as suggested by the cylinder climbing assay. No differences were observed in the average time taken to cover a track or total distance climbed during the duration of the assay (Fig. 4E,F). Taken together, these assays suggest that ELS flies exhibit motor deficits. It remains to be discovered if these are originating in the muscle or nervous system or is a combination of aberration in both. It must be recalled that locomotory circuits and muscle architecture for climbing as well as flight are significantly remodeled during pupal to adult development, as larva lack legs and wings.

## Discussion

*Drosophila melanogaster* as a model animal offers an excellent opportunity to explore how nutritional stress during development impacts adult phenotypes. We used the system to ask how late larval starvation manifests in adults on a normal diet. Consistent with literature, reduced food availability during development results in adult ELS (early life starved) flies that are smaller (Fig. 1D,E). In terms of systemic biochemical readouts, when normalized to body weight, there were no significant changes observed in overall protein, triacylglyceride (TAG), glycogen or trehalose levels, except for lower glucose levels specifically in males (Fig. 2A-E). Measurements in the hemolymph revealed higher TAGs in ELS females and lower trehalose in ELS males (Fig 2G,I). These results do not immediately point towards an obvious metabolic defect. It would appear that in steady-state, normal diet conditions, ELS females have altered lipid homeostasis (Fig. 2B and 2G) and ELS males have altered sugar homeostasis (Fig. 2 D,E, H and I). Further studies are required to uncover why this is so.

A few gut-related parameters were investigated in female ELS flies, of which two phenotypes – increased proportion of flies with higher gut barrier permeability (Fig. 3A) and slower activation of antimicrobial response to enteric infection (Fig 3E,F) – were significantly different. It would be interesting to understand the implication of these findings on adult gut function and immune response.

*Drosophila* has a large repertoire of behavioural assays of which a very small subset was attempted with ELS flies. Both climbing (Fig. 4A) as well as air-puff stimulated tethered flight (Fig. 4B) were different from controls. While climbing was lower in males and females, flight duration was lower for ELS females and higher for ELS males. High-speed tracking reveled higher climbing speed of ELS females (Fig. 4C) and poorer geotactic index of ELS males (Fig. 4D). Further studies are required to understand if these differences are neuronal or muscular in origin. It would also be interesting to understand if higher-level cognitive tasks such as egg-laying decision and memory are changed in ELS adults. Finally, these studies reveal that nutritional insults during development impact adults in a sex-dependent manner, to the extent that in some phenotypes, such as flight behaviour (Fig. 4B), the differences occur in opposite directions.

To experimentally study how nutrition during development impacts adult phenotypes, several variables can be manipulated. Some of these are: developmental time at which nutritional deprivation is imposed, duration of nutritional deprivation, type of nutritional deprivation (complete, diluted or specific macronutrient based), type of diet fed to early life malnourished adults and the adult age at which phenotypes are measured. Given our desire to model public health nutrition paradigms in flies, we chose to understand the long-term implications of severe childhood wasting which is addressed by institutional feeding. The WHO classifies severe wasting in infants or children < 5years when their weight for height are more than 3 standard deviations from normal (https://www.who.int/health-topics/malnutrition). Once identified, these children are placed on therapeutic foods for recovery. What happens specifically to this cohort of individuals as adults is not documented but given the timeframe of when they reach adulthood, with increased economic prosperity, some of them would be consuming a normal diet. Hence, we adopted an experimental paradigm of acute larval starvation followed by a “compensatory” normal adult diet. Incidentally, epidemiological studies point to a correlation between early life malnourishment and increased risk of cardiovascular disease and metabolic risk in adults (Wells *et al*., 2019), which is conceptualized as DoHAD (developmental origins of adult health and disease). Interestingly, when exposed to a high-fat diet, both male and female ELS adults displayed enhanced accumulation of TAGs as compared to their respective controls (Fig 2J), which provides support to use *Drosophlia* as a model to study the DoHAD concept.

Although this article describes a small subset of phenotypes, the results underline the importance of correct and timely nutritional inputs during development. Several avenues of investigations can be pursued hereafter. First, the causal mechanisms that drive metabolic, gut and behavioural phenotypes can be studied. It would be interesting to learn how nutritional stress “re-programs” various classes of tissues using the *Drosophila* genetic and behavioural toolkit. Second, it opens up a window of opportunity to study how diet therapies can improve or modify adult phenotypes. For instance, gut permeability and immune response in ELS flies was altered. We propose that our experimental protocol and the observed gut phenotypes can be used to understand what diets, nutritional supplements or natural plant products can help improve gut barrier function and immune responses in syndromes such as EED. Third, our data suggests that the manifestation of larval malnutrition in adults is sex-dependent. This requires further interrogation because from a public policy perspective, gender is typically not considered while evaluating or planning public health nutrition campaigns in malnourished infants and children. It must be acknowledged that there are significant differences in fly and human development. Further, it is not possible to model all other environmental stressors such as post-natal care, pollution, hygiene or climatic circumstances, that strongly modify human development. Thus, our model is a biological approximation and cannot directly address the real-world scenario. Nonetheless, our results point to the exciting utility of using *Drosophila* to understand how early life nutritional inputs continue to determine adult functions.

## Acknowledgements

This work started while M was a postdoc at NCBS-TIFR with an India Alliance Early Career Fellowship #IA/E/12/1/500742. We are deeply grateful to Gaiti Hasan for lab space and mentorship. This work also was partly supported by funding from Bill and Melinda Gates Foundation Sentinel grant to M (BIRAC/PMU/2019/Sentinels-005). Salary and small equipment support to M from a grant by Rural India Supporting Trust (RIST) to TDU is acknowledged. Thanks to Sveta Chakraborti (IISc) for providing the *Ecc15* strain.

## Author Contributions

Investigations were performed by SGP, SS, AA, OTS and DR. Conceptualization, formal analysis, funding acquisition, supervision and writing – review & editing by Megha.

